# A nonsense *mutation in Myelin Protein Zero* causes congenital hypomyelination neuropathy through altered P0 membrane targeting and gain of abnormal function

**DOI:** 10.1101/352112

**Authors:** Pietro Fratta, Francesca Ornaghi, Gabriele Dati, Desirée Zambroni, Paola Saveri, Sophie Belin, Patrizia D’Adamo, Michael Shy, Angelo Quattrini, M. Laura Feltri, Lawrence Wrabetz

**Author notes:** These authors contributed equally to this work (Pietro Fratta, Francesca Ornaghi, Gabriele Dati).

## Abstract

Protein Zero (P0) is the major structural protein in peripheral myelin and mutations in the *Myelin Protein Zero* (*Mpz*) gene produce wide ranging hereditary neuropathy phenotypes. To gain insight in the mechanisms underlying a particularly severe form, congenital hypomyelination (CH), we targeted mouse *Mpz* to encode P0Q215X, a nonsense mutation associated with the disease, that we show escapes nonsense mediated decay and is expressed in CH patient nerves. The knock-in mice express low levels of the resulting truncated protein, producing a milder phenotype when compared to patients, allowing to dissect the subtle pathogenic mechanisms occurring in otherwise very compromised peripheral myelin. We find that P0Q215X does not elicit an unfolded protein response, which is a key mechanism for other pathogenic *MPZ* mutations, but is instead aberrantly trafficked to non-myelin plasma membranes and induces defects in radial sorting of axons by Schwann cells (SC). We show that the loss of the C-terminal YAML motif is responsible for P0 mislocalisation, as its addition is able to restore correct P0Q215X trafficking *in vitro*. Lastly, we show that P0Q215X acts through dose-dependent gain of abnormal function, as wildtype P0 is unable to rescue the hypomyelination phenotype. Collectively, these data indicate that alterations at the premyelinating stage, linked to altered targeting of P0, may be responsible for CH, and that different types of gain of abnormal function produce the diverse neuropathy phenotypes associated with *MPZ*, supporting future allele-specific therapeutic silencing strategies.

## Introduction

Mutations in the *myelin protein zero* gene (*MPZ*) are a major cause of inherited neuropathy and can manifest in a wide range of clinical phenotypes, ranging from early onset forms, as Dejerine-Sottas syndrome (DSS) and Congenital Hypomyelination (CH), to less severe, adult onset, Charcot-Marie-Tooth diseases (CMT) (Warner et al, 1996).

Protein Zero (P0), encoded by the *MPZ* gene, is primarily expressed in Schwann cells (SC) and is the major structural transmembrane protein in peripheral myelin. P0 plays a crucial role in myelin formation by compacting adjacent wraps of SC plasma membrane, through *in trans* homophilic adhesion of its extracellular domain (D’Urso et al, 1990; Shapiro et al, 1996; Greenfield et al, 1973; Filbin et al, 1990). To date there are over 100 mutations in *MPZ*, known to cause peripheral neuropathies in patients and, although most of them are located in the extracellular domain of P0, few mutations in the short cytoplasmic domain were also reported (Warner et al, 1996; Baets et al, 2011). The intracellular tail of P0 has several putative functions, including compacting the cytoplasmic apposition in myelin (Martini et al, 1995), modulating adhesion by the P0 ectodomain (Xu et al, 2001; Wong et al, 1994 and 1996) and directing trafficking of P0 through the YAML motif (Kidd et al, 2006).

We have previously shown that different *MPZ* mutations act through distinct pathogenic mechanisms (Wrabetz et al, 2006; Pennuto et al, 2008). To further our understanding of how *MPZ* mutations cause disease, we generated a mouse model carrying the *MPZ*Q215X mutation which should delete part of the cytoplasmic domain and causes the severe CH neuropathy (Warner et al, 1996; Mandich et al, 1999; Shy et al, 2004). Here we show that lack of the last 33 aminoacids induces an altered trafficking of the protein to non-myelin plasma membranes and is associated with altered radial axonal sorting by SCs in the early phases of myelination. We demonstrate that *Mpz* Q215X acts through a dose-dependent, gain of abnormal function which cannot be rescued by supplementing normal P0, carrying important implications for therapeutic strategies.

## Materials and methods

All materials and methods can be found in the Supplementary material.

## Results

### Q215X escapes NMD and Mpz^Q215X/+^ mice express low levels of mutant P0

The CH-causative *MPZ* Q215X mutation induces a premature stop codon and, due to its location in the penultimate exon, has been predicted to escape nonsense mediated decay (NMD) (Inoue et al, 2002). We therefore first tested the expression of the mutant and WT *MPZ* alleles from a skin biopsy of a patient carrying this mutation and found equal expression levels of the two alleles, supporting the prediction that the mutation is not subject to NMD (**Fig. 1A**). To allow disease mechanism investigations, we generated knock-in mouse lines by homologous recombination in ES cells (**Fig. S1**). We generated ES cells carrying the Q215X mutation and two control lines (LoxPA3 and LoxPD1), where we inserted the WT sequence. Q215X heterozygous (*Mpz^Q215X/+^*) and homozygous (*Mpz^Q215X/Q215X^*) mice along with LoxP controls (*Mpz^LoxP/+^* and *Mpz^LoxP/LoxP^*) were all viable. The Q215X mutation is predicted to encode for a truncated P0, lacking 33 amino acids of its C-terminal intracellular domain. Western blot analysis on sciatic nerve (SN) protein extracts of P28 wild type, *Mpz^Q215X/+^* and *Mpz^Q215X/Q215X^* mice confirmed the mutation produces a smaller P0 with a molecular weight of approximately 24kD (**Fig. 1B, F**). Surprisingly, the amount of the mutated protein was strongly reduced relative to endogenous P0. We therefore tested the levels of *Mpz* RNA observed a reduction to 25% of WT levels (**Fig. 1D, E**). The mouse Q215X transcript was not targeted by NMD (data not shown), as predicted from patient data above, and the reduction in expression was present also in control lines carrying only the LoxP site (**Fig. 1D, E**). Importantly, although the presence of LoxP accounts for a reduction at the RNA level, the amount of truncated protein was reduced in Q215X homozygous nerves compared to equally expressing LoxP homozygous nerves, suggesting that the mutation affects the protein stability (**Fig. 1F, G**). Thus, the Q215X knock-in mouse generated a truncated protein that was expressed at low levels, probably due to the combined effect of the LoxP site at the transcriptional level and of the truncation at the protein level.

**Figure 1.**
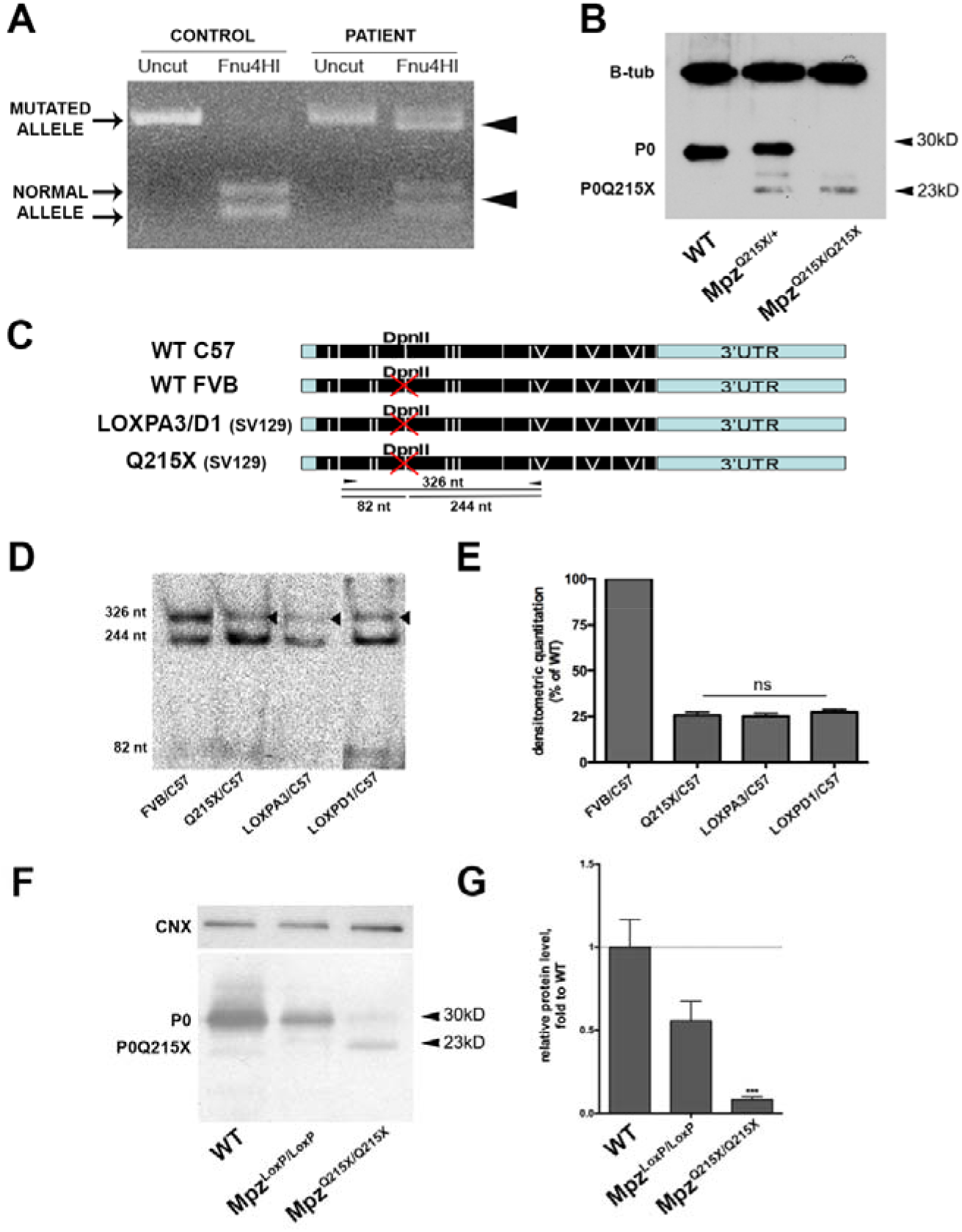
Q215X escapes NMD and Mpz^Q215X/+^ mice express low levels of truncated protein. **A.** Retrotranscribed and amplified *MPZ* RNA from a skin biopsy of a patient carrying the Q215X mutation shows similar expression of the mutated (uncut with Fnu4HI) and wildtype (cut) alleles. As expected all transcript is wildtype (cut) in a control sample. **B.** Western blot using anti-P0 and anti-βTubulin on protein extracts from sciatic nerves of *Mpz*^+/+^, *Mpz^Q215X/+^* and *Mpz^Q215X/Q215X^* sacrificed at postnatal day 28 show the presence of a truncated form of the protein at 23kD **C.** Schematic representation of the DpnII restriction enzyme sites in the different *Mpz* alleles. **D.** DpnII restriction enzyme digestion of the RT-PCR-amplified cDNA, obtained from RNA extracted from *Mpz*^+/+^ (FVB/C57Bl6), *Mpz^Q215X/+^* (129SVPas/C57Bl6) and two independent lines of *Mpz^LoxP/+^* (129SVPas/C57Bl6) P28 sciatic nerves. Given all targeted vectors were generated on 129SVPas background, and therefore did not carry the DpnII restriction site, we crossed all to C57Bl6 background, where the wt Mpz allele does have a DpnII restriction site. This allowed differentiation of WT *Mpz* (cut) and targeted alleles (uncut) by digestion. A WT mouse carrying one FVB allele (uncut) and C57Bl6 allele (cut) was used as technical control and showed equal amounts of expression from the two alleles. **E.** Quantification of uncut band vs digested product (FVB WT transcript) are plotted showing a similar reduction in all samples carrying the LoxP site (n=3 mice/genotype; oneway ANOVA, Tukey’s post-test, p value *ns* between samples carrying the LoxP site. Error bars represent SEM). **F.** anti-P0 western blot of protein extracts from *Mpz*^+/+^, *Mpz^LoxP/LoxP^* and *Mpz^Q215X/Q215X^* p28 sciatic nerves. **G.** Quantification of P0 normalised to calnexin is shown (n=3 mice/genotype; 2way ANOVA, Bonferroni post-test, p value<0.001 *vs* WT. Error bars represent SEM).

### Mpz^Q215X/+^ mice develop a neuropathy with radial sorting defects

Whilst the Q215X mutation in patients induces an early disease onset and severe clinical features, the *Mpz^Q215X/+^*mice showed no gross phenotype compared to controls and did not reveal any external sign of peripheral neuropathy (gait difficulty, tremor, atrophy of the hindlimb musculature). In order to assess whether the mutant P0Q215X induces nerve abnormalities, we analysed semithin sections (STS) of P28 sciatic nerves from *Mpz^Q215X/+^* mice, which revealed a mild hypomyelination when compared to controls (**Fig. 2A**). G-ratio analysis to assess myelin thickness, confirmed the hypomyelination in *Mpz^Q215X/+^* and showed that was similar to the hypomyelination seen in P0 heterozygous knockout (*Mpz^+/−^*) mice, which served as control for reduced *Mpz* expression (G-ratio Q215X/+ 0.69, P0+/− 0.7, WT 0.65; **Fig. 2B**).

**Figure 2.**
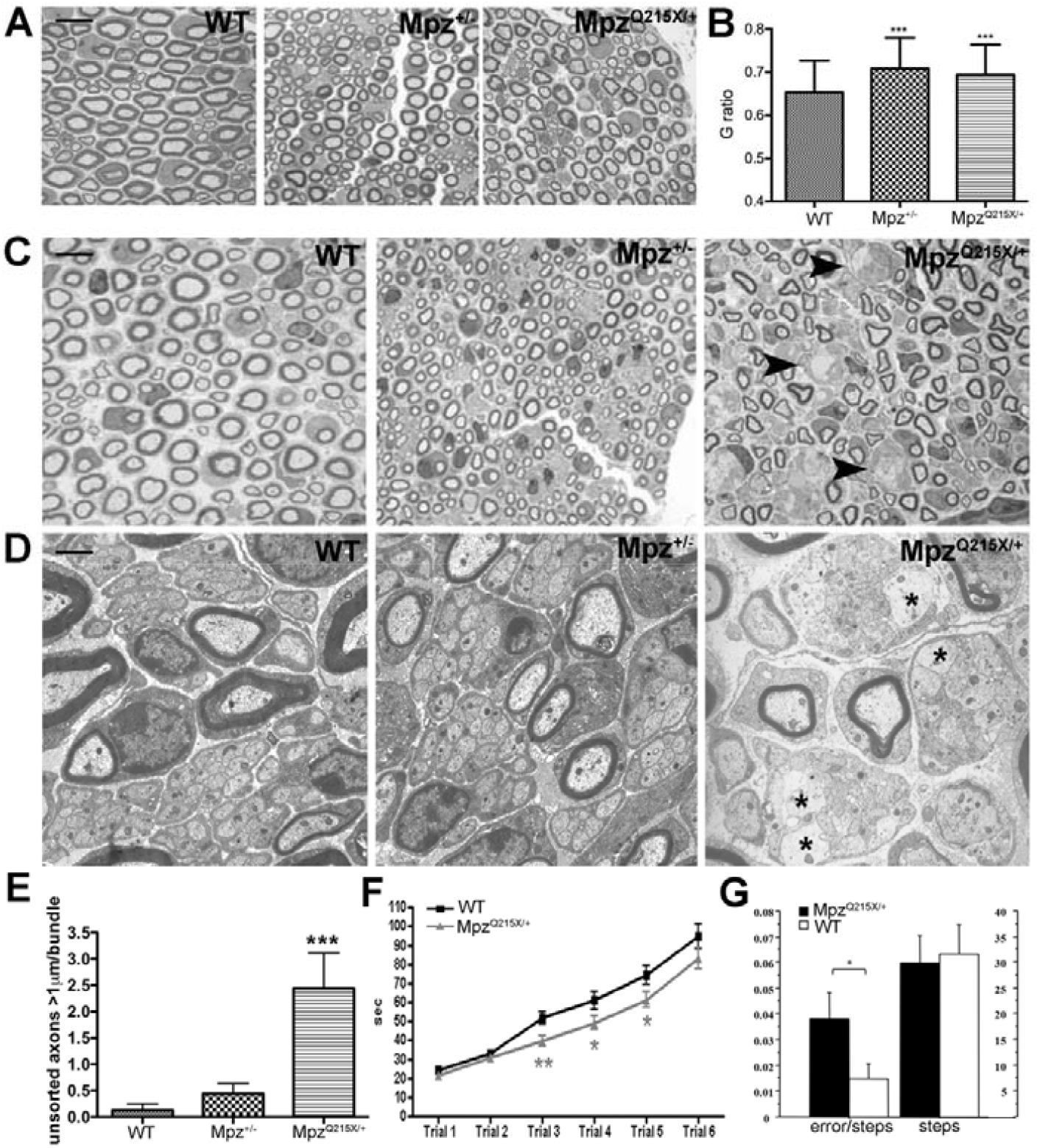
P0Q215X induces radial axonal sorting defects and neuropathy. **A.** Semithin Sections (STS) of p28 sciatic nerves taken from *Mpz*^+/+^, *Mpz^+/−^* and *Mpz^Q215X/+^* mice. Magnification 100X, scale bar 10μm. **B.** G ratio quantifications of (A) (g ratio *Mpz*^+/+^ 0.65+/−0.002; g ratio *Mpz^+/−^* 0.7+/−0.002; g ratio *Mpz^Q215X/+^* 0.69+/−0.002); n=3 mice/genotype, paired t-test, p value<0.001 *vs* WT. Error bars represent SEM). **C.** STS of p11 sciatic nerves from *Mpz*^+/+^, *Mpz^+/−^* and *Mpz^Q215X/+^* mice mice. Arrowheads indicate bundles of unsorted axons. **D.** Electromicrographs of p11 sciatic nerves taken from *Mpz*^+/+^, *Mpz^+/−^* and *Mpz^Q215X/+^* mice, showing details of bundles of unsorted axons (magnification 20,000x; scale bar 1μm). **E.** Number of unsorted axons (in direct contact with other axons, indicated by asterisks in the right panel) within the bundles, with a diameter > 1μm (n=3 mice/genotype, one-way ANOVA, Tukey test, p<0.001 *vs* WT. Error bars represent SEM). **F**. Rotarod analysis shows that *Mpz^Q215X/+^* mice at P11 (n=72) remain on the accelerating cylinder for significantly less time than wild type littermates (n=57). (paired t-test, *p<0.05 and **p<0.01 *vs* WT. Error bars represent SEM). **G**. Grid walking test shows that *Mpz^Q215X/+^* mice at P15 (n=32) had a number of errors as compared to wild to littermate controls (*Mpz*^+/+^; n=35) with the same number of total steps. (ANOVA genotype effect for errors F[1,65]=8.33; p=0.005. Error bars represent SEM).

In patients the Q215X mutation causes a deficit during development, we thus focused our analysis in the first two weeks of postnatal life. At P11, we observed bundles of unsorted mixed caliber axons in *Mpz^Q215X/+^* nerves (**Fig. 2C**, arrowheads) and not in wild type littermates or in *Mpz^+/−^* mice (**Fig. 2C**). Unsorted axons were significantly augmented in *Mpz^Q215X/+^* compared to both WT and *Mpz^+/−^* (**Fig. 2D, E**), and at P14, the radial sorting defect was not visible anymore (data not shown) indicating that the deficit in P11 *Mpz^Q215X/+^* mice is that of a delay in myelination. We tested whether this defect resulted in a neuromuscular deficit using the rotarod and grid walking tests. *Mpz^Q215X/+^* P10 to P12 mice showed a significant reduction in motor coordination compared to controls, revealing that the transient defect in the process of myelination identified in P11 mice results in a developmental motor deficit (**Fig. 2F**). A similar impairment was also detected using the grid walking test where P15 *Mpz^Q215X/+^* mice had a significantly higher number of errors compared to controls (ANOVA genotype effect F[1,65]=8.33; p=0.005, normalized on the same number of steps: ANOVA genotype effect F[1,65]=0.4; p=0.52) (**Fig. 2G**). These neuromuscular deficits are not present anymore in adult (P28) mice (data not shown). Taken together, these data show that the Q215X mutation, even expressed at low levels, induces an axonal radial sorting defect with delay in myelination leading to a dysmyelinating neuropathy. This early defect in postnatal life parallels the dysmyelinating disease reported for the two patients presenting Q215X de novo mutations and is specifically due to the presence of the Q215X mutated glycoprotein, and not to the lower expression of *Mpz*, as demonstrated by the absence of unsorted bundles of large calibre axons in P0+/− mice. These observations collectively suggest that Q215X acts via a gain of function (GOF) mechanism.

### P0Q215X does not induce ER stress and reaches the plasma membrane

Altered intracellular retention inducing ER stress has been previously associated and shown to play a critical role in the pathogenesis of gain of function *MPZ* mutations (Wrabetz et al, 2006; Pennuto et al, 2008; Saporta et al., 2012), prompting us to investigate whether P0Q215X induces ER stress and an unfolded protein response.

First, we generated *Mpz^Q215X/Q215X^*, that only express the mutant protein, and determined whether P0Q215X was retained in the ER by performing co-staining with the ER marker KDEL on teased nerve fibers. We compared *Mpz^Q215X/Q215X^* to WT (**Fig. 3A**) and also to LoxP homozygous (*Mpz^LoxP/LoxP^*) mice (**Fig. 3B**), which have a similar amount of *Mpz* expression. We could not detect P0Q215X in the ER (**Fig. 3C**), differently from P0S63del that we used as a positive control (**Fig. 3D**). Further, mRNA levels of ER stress transcripts *BiP* and spliced *Xbp-1*, were not significantly increased in *Mpz^Q215X/Q215X^* nerves, in contrast to nerves from mice carrying the P0S63del mutation, showing that the Q215X mutation does not induce ER stress (**Fig. 3E**). P0Q215X was detected in the myelin of teased nerve fibers (**Fig 3C**), suggesting it can be normally trafficked. P0 forms tetramers before being targeted to myelin, and we tested whether P0Q215X is able to interact with its wildtype (WT) counterpart. We generated *Mpz^Q215X/Q215X^* mice expressing a myc-tagged P0WT (P0myc), allowing to perform an immunoprecipitation assay using an anti-myc antibody, in this setting specific to P0WT (Fratta et al, 2011). Western blot analysis of immunoprecipitates demonstrated pulldown of P0Q215X, supporting the interaction between P0Q215X and WTP0 (**Fig. S2**). Overall, this data shows that P0Q215X reach the plasma membrane, does not induce ER stress and can interact with P0WT.

**Figure 3.**
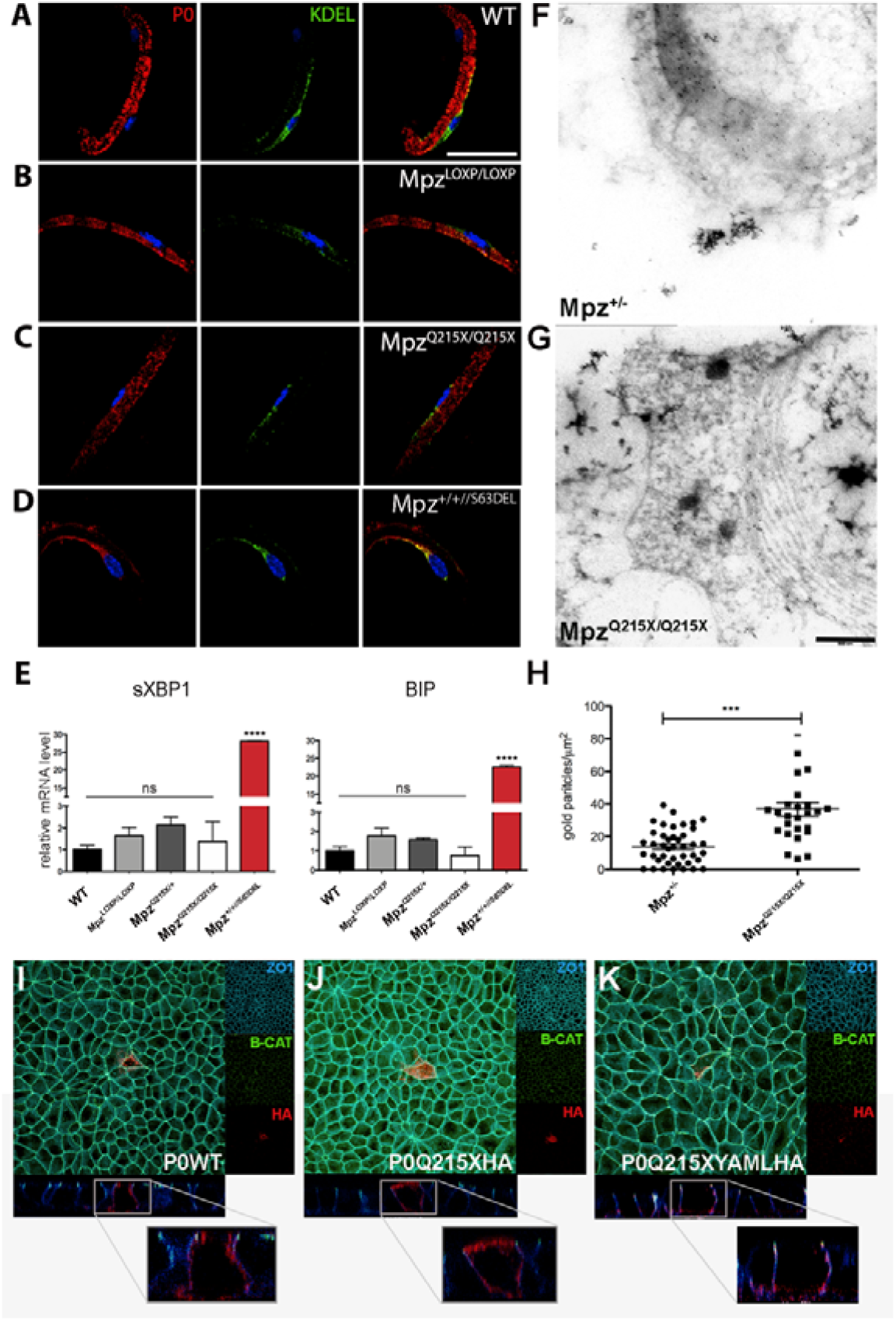
Lack of the YAML motif induces P0Q215X mis-trafficking to non-myelin membrane. Immunohistochemistry on teased sciatic nerve fibers from *Mpz^Q215X/Q215X^* (**C**) and *Mpz^LoxP/LoxP^* (**B**) mice show P0 ER retention in Mpz^+/+//P063del^ mice (**D**) but not in *Mpz^Q215X/Q215X^* and WT controls (**A**). DAPI (blue); P0 (red); KDEL (green). Scale bar: 35 μm **E.** qPCR quantifications of XBP-1 and BiP levels show no significant changes in *Mpz^Q215X/Q215X^* (n=3 mice/genotype; Oneway ANOVA, Tukey’s post-test. P value vs WT ns and <0.001 (****). Error bars represent SEM). **F, G.** ImmunoEM using anti-P0 Ab on *Mpz^Q215X/Q215X^* and *Mpz^+/−^* p10 sciatic nerves (scale bar 500 nm). **H.** Quantification of gold granules counted in non-myelin plasma membranes shows a statistically significant difference between *Mpz^Q215X/Q215X^* and *Mpz^+/−^* (n=3 mice/genotype paired; t-test, p value <0.0001; error bars represent SEM). Transiently transfected MDCK cells expressing P0Q215X-YAML-HA, or P0Q215X-HA and P0WT-HA as controls, were stained for HA (red), ZO1 (blue, marker of basolateral surface) and Beta Catenin (green, marker of apical surface). Three independent replicates. **I.** P0WT-HA is targeted to the basolateral surface of MDCK cells: in XZ confocal images, P0 immunostaining did not colocalize with apical marker Beta Catenin but was concentrated in basolateral membranes, which are immunostained for ZO1. **J.** P0Q215X-HA was targeted to the apical surface and/or accumulated throughout the cell. **K.** P0Q215X-YAML-HA is localized to the basolateral surface, similarly to P0WT-HA.

### P0Q215X is incorrectly trafficked to non-myelin plasma membranes

As the C-terminal domain has been shown to be important in targeting P0 to the membrane, we performed immunoelectronmicroscopy (IEM) on p10 *Mpz^Q215X/Q215X^* and *Mpz^+/−^* mice to assess this process. Gold granules indicate that P0 molecules are located primarily in the myelin sheath in *Mpz^+/−^* nerves (**Fig. 3F, G**), but interestingly, gold granules were frequently detected outside of myelin in *Mpz^Q215X/Q215X^* nerves, suggesting that P0Q215X could be partially mis-trafficked to non-myelin plasma membranes. Quantification of P0 showed an increase of the number of gold granules in non-myelin plasma membranes from *Mpz^Q215X/Q215X^* nerves, which is significantly higher than in the *Mpz^+/−^* controls (**Fig. 3H**). These data demonstrate that P0Q215X trafficking is altered *in vivo* with P0 being targeted to non-compact myelin membranes.

### The YAML sequence in the cytoplasmic tail regulates P0 trafficking

P0Q215X lacks a YAML sequence in its C-terminal region, which has previously been implicated in myelin membrane targeting (Kidd et al, 2006). In order to address whether this determines the altered trafficking of P0Q215X, we used an *in vitro* assay where the YAML motif directs P0 trafficking to the basolateral surface of transfected and polarized MDCK cells (Kidd et al, 2006). We transfected MDCK cells with P0Q215X-HA or P0Q215X-YAML-HA, and P0wt-HA as control, and stained the cells for HA, for Beta-catenin, a marker of the apical junction, and for ZO1, a marker of the basolateral surface. P0WT-HA was present at the basolateral surface of the cells (**Fig. 3I**), whereas the mutant P0Q215X-HA did not respect the basolateral distribution and was detected also at the apical surface (**Fig. 3J**). In contrast, the P0Q215X-YAML-HA construct was detected only at the basolateral surface of MDCK cells (**Fig. 3K**). These data show that YAML is sufficient to restore proper surface membrane targeting of P0Q215X, suggesting its deficiency determines the observed altered targeting of P0Q215X.

### P0Q215X acts through a gain of abnormal function

Whether mutations act though gain or loss of function is a crucial information as it has implications for therapeutic approaches. We reasoned that if the phenotype observed in *Mpz^Q215X/Q215X^* mice was due to hypomorphic P0 expression, i.e. a loss of function, then *Mpz^LoxP/LoxP^* should have a similar phenotype, whilst if GOF were involved, the phenotype should be aggravated in a dose-dependent manner. To therefore test whether Q215X induces disease by a dosage-dependent GOF, we first generated *Mpz^Q215X/Q215X^* mice and we compared myelination to that of *Mpz^LoxP/LoxP^* mice, that express equivalent levels of wt *Mpz.* Whilst the reduced amount of WT *Mpz* expressed in *Mpz^LoxP/LoxP^* mice (∼20% of normal) was sufficient for myelination in SNs at P10, *Mpz^Q215X/Q215X^* animals (also expressing ∼20% of normal) manifested a very severe hypomyelination at this age (**Fig. 4A**). To further eliminate the confounding element of LOF in *Mpz^Q215X/Q215X^* mice, we crossed *Mpz^Q215X/Q215X^* mice with a P0 overexpressor transgenic mouse line (*Mpz^+/+;tgP0oe^*) (Wrabetz et al, 2000). The *Mpz^+/+;tgP0oe^* mice express *Mpz* wt at 80% on top of normal (where normal refers to two *Mpz* wt alleles) and were previously demonstrated to fully rescue the P0 null phenotype at P28 (Wrabetz et al, 2000). We compared *Mpz^Q215X/Q215X;tgP0oe^* mice to *Mpz^−/−;tgP0oe^* mice at P10. Consistent with previous results STS from *Mpz^−/−;tgP0oe^* sciatic nerves show normal myelination (**Fig. 4B**) with a G-ratio similar to wild type (**Fig. 4C**; G ratio values: WT 0.63; P0ko 0.76; P0ko//P0oe 0.63). Electron microscopic (EM) analysis confirmed the presence of normally compacted, thick myelin sheaths in these specimens (**Fig. 4D**) - demonstrating that the P0oe transgene is able to rescue the *Mpz^−/−^* phenotype also at P10. In contrast, STS from *Mpz^Q215X/Q215X;tgP0oe^* sciatic nerves showed abnormally thin myelin (**Fig. 4B**). EM analysis showed bundles of unsorted axons as previously described and, most remarkably, very severe hypomyelination in sciatic nerve (**Fig. 4D**). P0WT was unable to induce an amelioration when compared to *Mpz^Q215X/Q215X^*, and the G-ratio was not significantly different (**Fig. 4C**; G ratio values: Q215X/Q215X0.75; Q215X/Q215X//P0oe 0.74). These data conclusively demonstrate that Q215X causes the myelin phenotype by a dose-dependent GOF mechanism.

**Figure 4.**
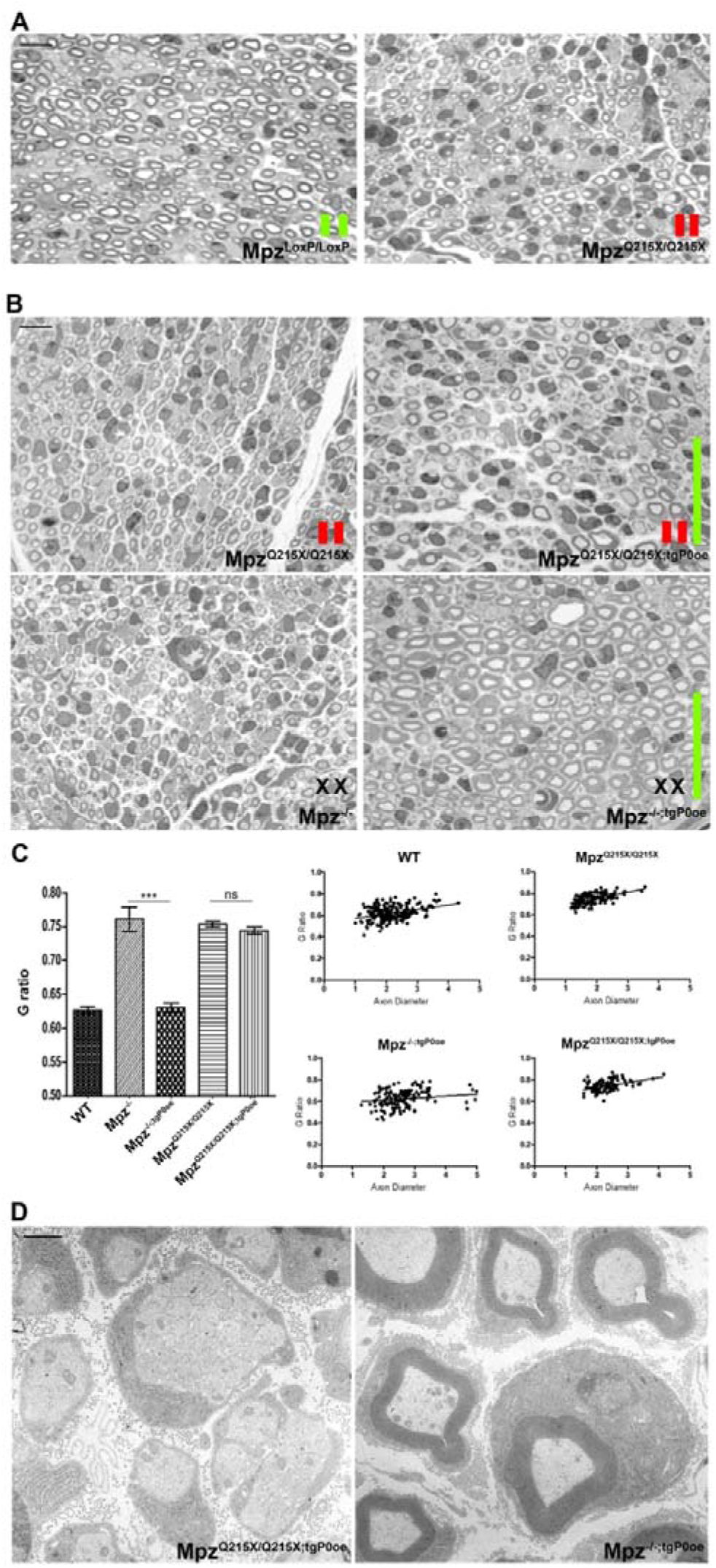
P0Q215X acts through gain of abnormal function. **A.** STS of sciatic nerves from P10 *Mpz^LoxP/LoxP^* and *Mpz^Q215X/Q215X^* mice. *Mpz^Q215X/Q215X^* nerves are hypomyelinated when compared to *Mpz^LoxP/LoxP^* nerves (scale bar 10μm). **B.** STS of P10 nerves from *Mpz^Q215X/Q215X^* (upper panel, at left) and from *Mpz^Q215X/Q215X^* mice overexpressing P0WT (*Mpz^Q215X/Q215X;tgP0oe^*, upper panel, at right); STS of P10 nerves from *Mpz*^−/−^ (lower panel, at left) and *Mpz*^−/−^ overexpressing P0WT (*Mpz^−/−;tgP0oe^*, lower panel, at right); scale bar 10μm. Restoration of normal levels of P0wt is unable to rescue the hypomyelination in *Mpz^Q215X/Q215X;tgP0oe^* (upper panel, at right) whereas it does rescue the very severe hypomyelination in *Mpz^−/−;tgP0oe^* (lower panel, at right). In A and B, the red and the green bars indicate the expression of the wild type (green) or mutated (red) alleles. **C.** G ratio measured at P10 shows hypomyelination in *Mpz^Q215X/Q215X;tgP0oe^*, whereas hypomyelination is rescued in *Mpz^−/−;tgP0oe^*, where the G ratio is comparable to WT mice. G ratio values: WT 0.63; *Mpz^−/−^* 0.76; *Mpz^−/−;tgP0oe^* 0.63; *Mpz^Q215X/Q215X^*0.75; *Mpz^Q215X/Q215X;tgP0oe^* 0.74. N=3 mice/genotype; paired t test: p<0.0001 *Mpz^−/−^ vs Mpz^−/−;tgP0oe^* and p value *ns Mpz^Q215X/Q215X^ vs Mpz^Q215X/Q215X;tgP0oe^*. Error bars represent SEM. **D.** Electron micrographs confirms the rescue of myelin thickness in *Mpz^−/−;tgP0oe^* at P10 (right image), whereas hypomyelination and the sorting defects still remain in *Mpz^Q215X/Q215X;tgP0oe^* (left image). Scale bar: 1μm.

## Discussion

We have generated a novel mouse model for congenital hypomyelination induced by a nonsense *MPZ* mutation. We first show, using patient tissue, that the mutation escapes nonsense mediated decay and is expressed similarly to its wild type counterpart. In both our mutant mice and in control lines, where a WT-Mpz allele was inserted and therefore carry a LoxP site, *Mpz* expression is reduced due to the presence of the LoxP sites in intron 5, and, as expected, the premature stop codon gives rise to a truncated protein in mice. This provides a model system where a very aggressive mutation is expressed at more limited levels, allowing us to work with a more tractable phenotype. Interestingly, protein levels appear lower compared to wild type LoxP alleles, suggesting reduced protein stability. The mutant heterozygous mice, in keeping with the reduced expression of mutant *Mpz*, develop a phenotype which is less severe than that of patients, but nonetheless show hypomyelination with reduced nerve conduction velocity. We were also able to identify a novel defect in axonal radial sorting during myelination at P10, which is intriguing, as defects at the axonal sorting stage are compatible with the early-onset developmental phenotype observed in patients.

Normally P0 is synthesized in the ER, glycosylated, assembled as tetramers and then targeted to the myelin membrane, and early stages of this process have been shown to be altered in other disease-causing mutations (Wrabetz et al, 2006; Pennuto et al, 2008; Saporta et al, 2012). We therefore used a combination of molecular mouse tools, biochemical, confocal microscopy and immunoEM approaches to investigate P0Q215X maturation, and show that P0Q215X is able to correctly mature, interact with its WT counterpart and be trafficked to the membrane. No ER stress is induced, clearly differentiating the mechanism of action from other *MPZ* mutations causative of CMT or CH that we previously characterized in mice (Wrabetz et al, 2006; Pennuto et al, 2008; Saporta et al, 2012).

As one of the known functions for P0 cytoplasmic domain, which is truncated in P0Q215X, is to target the protein to myelin (Kidd et al, 2006), we analysed in detail the membrane targeting of the protein, and found that Q215X was increasingly mis-directed to non-myelin plasma membranes. We then used an *in vitro* assay to show that re-insertion of the YAML in the cytoplasmic domain is able to rescue the trafficking of P0Q215X. Interestingly, even wild type P0, if overexpressed at very high levels, is mislocalized at the abaxonal membrane and mesaxon, causing Congenital Hypomyelination due to radial sorting defects and amyelination. The latter is probably due to a gain of normal P0 function that causes homophilic P0 adhesion between apposed mesaxonal membranes (Wrabetz et al, 2000; Yin et al, 2000). These findings have important implications as potentially the targeting of an adhesive protein such as P0 to incorrect membrane domains may contribute also to the altered axonal radial sorting process, where SC membranes need to extend between and isolate axons.

The mis-targeting of P0 protein is compatible in principle with both gain and loss of function mechanisms. Dissecting the contribution of gain and loss of function is extremely important, especially in light of the recent increase in genetic therapy approaches, where for example PMP22 is targeted for reduction by means of antisense oligonucleotides in a CMT1A (Zhao et al, 2018). Our findings of an axonal sorting phenotype being present only in *Mpz^Q215X/+^* and not in *Mpz^+/−^* mice suggested a gain of function and in order to conclusively address this issue, we increased the dosage of mutant P0Q215X by breeding mice to homozygosity and then asked whether supplementation of P0WT protein would rescue the phenotype. Homozygous mice showed a very severe phenotype, but P0WT supplementation, which was able to rescue the full loss of P0 in *Mpz^−/−^* mice, was unable to correct the P0Q215X toxicity, strongly indicating GOF as the main mode of action.

The fact that different *MPZ* mutations have very distinct mechanisms of action, but the commonality of a GOF mechanism, has strong therapeutic implications. It makes it unlikely to develop a therapeutic approach targeting such a wide range of biological processes (e.g. altered membrane targeting, ER stress etc.), but supports a genetic therapeutic approach based on allele-specific knock down. Although currently gene silencing is being pursued only in conditions where the same mutation is present in a wide number of patients (e.g. CMT1A), future developments may use common polymorphism to develop an array of silencing tools. This will allow for a silencing approach to be applied to less frequent mutations such *MPZ*.

## Acknowledgements

We thank patients and families for their participation to research that has made this work possible. This work was supported by grants from the National Institute of Health (LW R01 NS55256 and R56NS096104), the Charcot Marie Tooth Association (LW) and Fondazione Telethon (to LW).

---

**SUPPLEMENTARY FIGURE 1.**
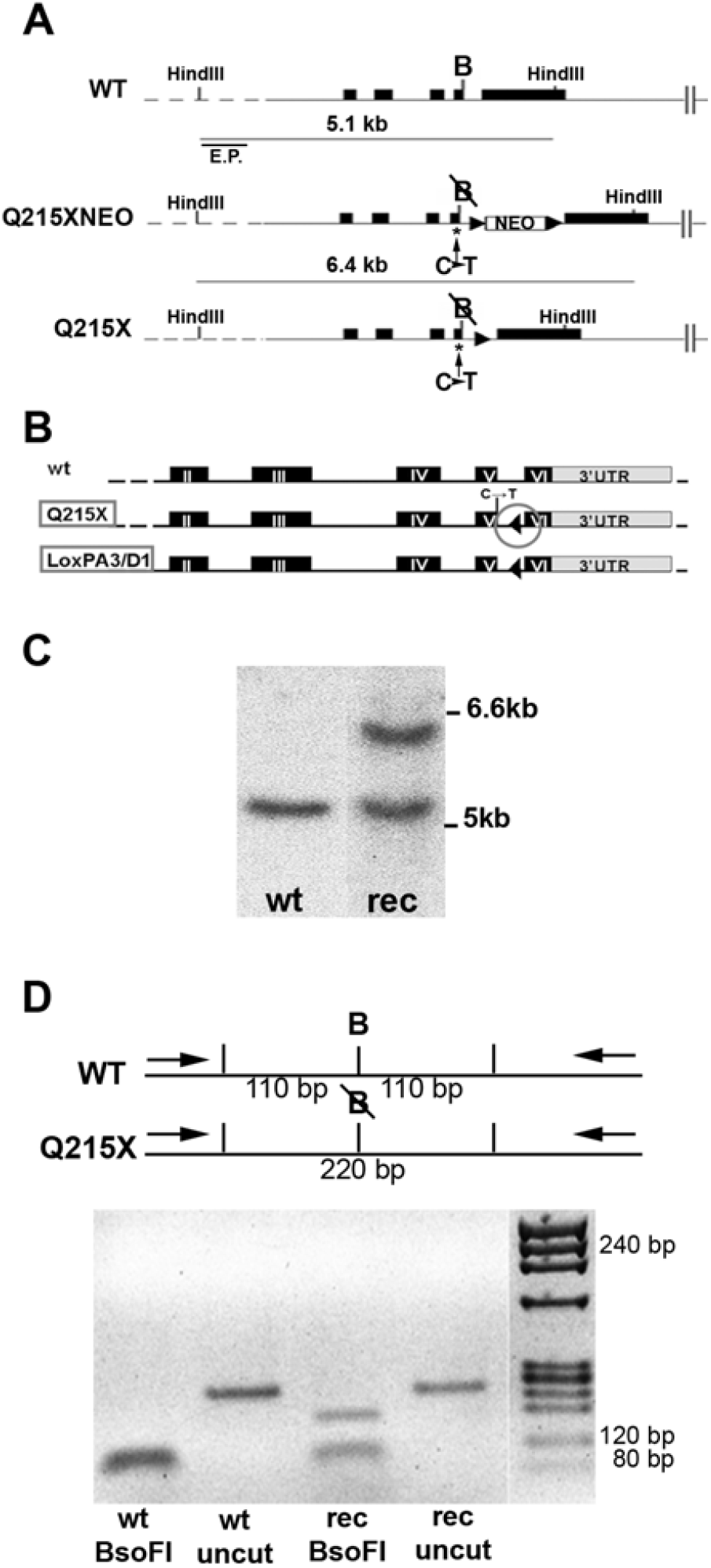
Generation of Q215X mouse line. **A.** Schematic representation of the genomic organization of wild-type, Q215XNEO and Q215X *Mpz* alleles. The external probe used for Southern blot analyses of ES cell clones is indicated (E.P.), together with the length of the DNA fragments, originating upon HindIII digestion. The C to T mutation in exon 5 is indicated by asterisks. The BsoFI site within exon 5 is indicated (B). **B.** Structure of P0Q215X and LoxP gene with the LoxP site (circle) in the intron 5. **C.** Southern Blot analysis of the genomic DNA of the ES cell clone where homologous recombination occurred. **D.** BsoFI restriction enzyme digestion of the PCR-amplified genomic region flanking the C to T mutation in exon 5. The BsoFI site within exon 5 is indicated (B).

**SUPPLEMENTARY FIGURE 2.**
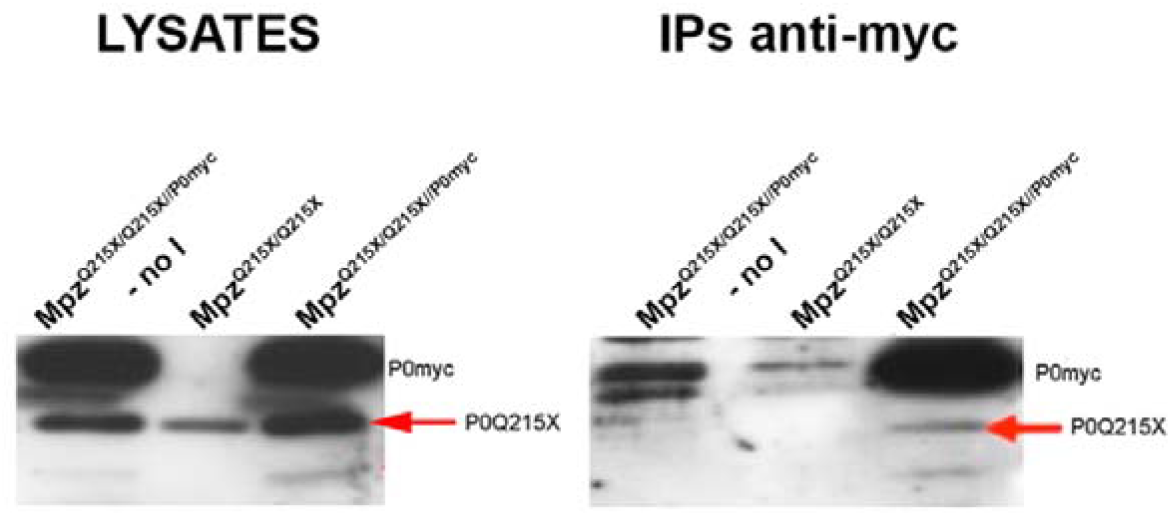
Interaction between P0Q215X and WTP0. Immunoprecipitation for P0myc in *Mpz^Q215X/Q215X^*^//P0myc^ mice. The red arrow indicates the bands corresponding to the truncated P0Q215X in the lysates and in the IP. Anti-myc antibodies are able to co-IP P0myc and P0Q215X from *Mpz^Q215X/Q215X^*^//P0myc^ mice. As negative controls, sciatic nerve lysates from *Mpz^Q215X/Q215X^*^//P0myc^ mice were exposed to only beads (no myc antibody, (no I)) or sciatic nerve lysates from *Mpz^Q215X/Q215X^* were immunoprecipitated with myc antibody. In these two lanes there are no bands corresponding to P0Q215X, demonstrating that P0Q215X is not simply adhering directly to the beads.

